# Development of the Pneumococcal Genome Library, a core genome multilocus sequence typing scheme, and a taxonomic life identification number barcoding system to investigate and define pneumococcal population structure

**DOI:** 10.1101/2023.12.19.571883

**Authors:** Melissa J. Jansen van Rensburg, Duncan J. Berger, Andy Fohrmann, James E. Bray, Keith A. Jolley, Martin C.J. Maiden, Angela B. Brueggemann

## Abstract

Investigating the genomic epidemiology of major bacterial pathogens is integral to understanding transmission, evolution, colonisation, disease, antimicrobial resistance, and vaccine impact. Furthermore, the recent accumulation of large numbers of whole genome sequences for many bacterial species enhances the development of robust genome-wide typing schemes to define the overall bacterial population structure and lineages within it. Using previously published data, we developed the Pneumococcal Genome Library (PGL), a curated dataset of 30,976 genomes and contextual data for carriage and disease pneumococci recovered between 1916-2018 in 82 countries. We leveraged the size and diversity of the PGL to develop a core genome multilocus sequence typing (cgMLST) scheme comprised of 1,222 loci. Finally, using multilevel single-linkage clustering, we stratified pneumococci into hierarchical clusters based on allelic similarity thresholds, and defined these with a taxonomic life identification number (LIN) barcoding system. The PGL, cgMLST scheme, and LIN barcodes represent a high-quality genomic resource and fine-scale clustering approaches for the analysis of pneumococcal populations, which support the genomic epidemiology and surveillance of this leading global pathogen.

**Impact statement:** Many thousands of pneumococcal genomes are available in the public domain, and this creates opportunities for the scientific community to re-use existing data; however, these data are most useful when the contextual data (provenance and phenotype) are also linked to the genomes. Therefore, we created a curated, open-access database in PubMLST that contained nearly 31,000 published pneumococcal genomes and the corresponding contextual data for each genome. This large and diverse pneumococcal database was used to create a novel cgMLST scheme and multilevel clustering method to define genetic lineages with high resolution and a standardised nomenclature. These are open-access resources for all to use and provide a unified framework for the characterisation of global pneumococcal populations.

## Introduction

In the first two decades of the 21^st^ century, the capacity to sequence the whole genome of microbes transformed the fields of microbiology and infectious disease. Furthermore, genomic epidemiology and surveillance are playing increasingly important roles in public health, vaccine development, and the assessment of vaccine impact [1–4]. Thousands of bacterial genome sequences are publicly available, and the potential for re-use of these genomes is of major benefit to the scientific community.

*Streptococcus pneumoniae* (the pneumococcus) is a bacterium that primarily resides in the healthy human nasopharynx but is also a major cause of localised and invasive infections worldwide. In 2019, prior to the COVID-19 pandemic, pneumococci were estimated to cause over 650,000 (95% UI 553,000–777,000) deaths due to pneumonia and nearly 45,000 (95% UI 34,700–59,800) deaths from meningitis among people of all ages [5]. Nonpharmaceutical interventions implemented during the pandemic to contain the spread of SARS-CoV-2 led to significant and sustained reductions in invasive pneumococcal disease (IPD), but IPD is returning to pre-pandemic levels in many countries [6–7]. Pneumococcal conjugate vaccines are used in many countries worldwide and have successfully reduced the global burden of pneumococcal disease over the past twenty years; however, the pandemic disrupted vaccination programmes worldwide and restoring these programmes is a public health priority [8].

As of 2023, there were tens of thousands of pneumococcal genome sequences in public repositories including the European Nucleotide Archive and GenBank; however, these repositories typically included minimal corresponding contextual data (provenance and laboratory data), and often the genomes were only available as unassembled short-read data, which limited their use by many investigators. Therefore, the first objective of this study was to create a Pneumococcal Genome Library (PGL) that would contain assembled genomes and corresponding contextual data from the peer-reviewed published literature and make those data freely available to all users via the PubMLST platform. We also incorporated basic genome quality criteria into the PGL to enable users, including non-specialists, to query, filter, re-analyse and download published pneumococcal genomes.

Originally, the pneumococcal population structure was defined using seven-locus multilocus sequence typing (MLST) and clonal complexes (CCs; clusters of related isolates), but recently, whole genome sequences have been used to define genetic lineages more precisely [9–12]. A well-defined population structure is the foundation on which other criteria can be mapped and interpreted, such as the lineages causing disease versus those found among healthy individuals, or antimicrobial-resistant lineages. Understanding the expected pneumococcal population structure is also necessary to detect and interpret any observed population-level changes to that structure. For example, pneumococci typically possess a polysaccharide capsule (serotype) and this is the primary vaccine antigen. There are over 100 antigenically different serotypes, and serotypes are typically associated with specific pneumococcal genotypes; therefore, new serotype/genotype combinations are a potential indication of capsular switching events that might have occurred in response to vaccination [13–15].

The MLST approach of defining alleles based upon sequence variation at a defined set of loci was transformative because sequence data are unambiguous and easily portable, and a common nomenclature was defined [16–17]. Assigning alleles and sequence types (STs) from large numbers of genomes is easily automated, whilst still relying on the expertise of a human data curator to ensure high quality data. Genomes can also be assessed using ribosomal MLST (rMLST), which characterises allelic diversity at the 53 bacterial ribosomal genes and is especially useful for species identification [18–19].

Core genome MLST (cgMLST) schemes have been implemented for several global pathogens to assess sequence variation at hundreds of core genes across the bacterial genome and increase the resolution of defined genetic lineages [20–25]. Therefore, the second objective of this study was to develop and implement a cgMLST scheme for pneumococci, which we extended to include a taxonomic life identification number (LIN) system to improve the resolution and clustering of genetic sublineages of pneumococci. The LIN approach was originally implemented for *Klebsiella pneumoniae* and has the main advantage of providing a definitive and stable ‘barcode’ for each genome that can be used to define and cluster groups of genetically related isolates, based on sequence variation within the core genes used in the cgMLST scheme [26–27].

Here, we describe the development of the PGL as a community resource of nearly 31,000 assembled pneumococcal genomes with their corresponding contextual data. We used the PGL to design a robust and reproducible pneumococcal cgMLST scheme of 1,222 core genes and developed a LIN barcoding system to cluster pneumococci across multiple levels, based upon sequence variation at those 1,222 genes. We implemented the PGL, cgMLST scheme, and LIN codes within PubMLST to provide open-access resources for the scientific community and a unified framework for the characterisation of global pneumococcal populations.

## Methods

### Literature search and article eligibility

We searched PubMed on 31 January 2020 using the terms “(‘Streptococcus pneumoniae’ OR ‘pneumococcus’) AND (‘genome sequencing’ OR ‘genome sequence’ OR ‘Genome, Bacterial’)”. An additional search was carried out on 15 July 2020 using the terms “(‘Streptococcus pneumoniae’ OR ‘pneumococcus’) AND ‘genome sequencing’”. Titles and abstracts of full-text, peer-reviewed articles published in English between 1 January 2000 and 15 July 2020 were manually screened for eligibility, and methods and results sections of an article were also reviewed if necessary. Eligible publications were original research articles that described genome data from at least one naturally occurring pneumococcal isolate.

### Genome data availability

The text and supplementary files of eligible published articles were searched for International Nucleotide Sequence Database Collaboration (INSDC) accession numbers, identifiers from other public databases, or genome data. For each article, we confirmed that the number of accessions, identifiers, or data files provided matched the total number of genomes described in the text, including any reference genomes and supplementary genome data. We also confirmed that any INSDC accession numbers were valid and corresponded to pneumococcal records.

### Data acquisition and genome assembly

Contextual data (country, year of isolation, source, diagnosis, sex, age, and serotype) were extracted from the published article and supplementary files, and data were manually cleaned to conform to allowed values accepted by the *S pneumoniae* PubMLST database. If available, assembled genomes were downloaded from the NCBI Nucleotide/Assembly databases. When assemblies were not available, short-read data was downloaded from ENA and genomes were assembled *de novo* using Velvet (v1.2.10) and VelvetOptimiser (v2.2.4) using a range of kmer sizes (19-191 bp) [28]. Assembled contigs shorter than 200 bp were removed. Genomes generated in earlier studies were assembled as previously described [9–10,29–40]. Illumina MiSeq data from two datasets assembled poorly with Velvet, so these genomes were assembled with SPAdes implemented in the INNUca pipeline v.4 [41–44]. Velvet assemblies were retained for the subset that did not assemble with the INNUca pipeline.

### Creation of the PubMLST Pneumococcal Genome Library

In the *S pneumoniae* PubMLST database, isolate records were created that included the assembled genome, any corresponding provenance or laboratory data, and the PubMed identifier of the original publication. A separate view of the *S pneumoniae* PubMLST database was created to enable users to access the PGL directly (https://pubmlst.org/organisms/streptococcus-pneumoniae/pgl). All known MLST and rMLST alleles were assigned by the BIGSdb autotagger tool in PubMLST [19]. New alleles and STs were manually curated and assigned by the database curators (ABB and JEB, respectively).

### rMLST species identification

The rMLST database (https://pubmlst.org/species-id) contains bacterial genomes compiled from the NCBI Assembly database and the assembly of short-read data [18–19]. The allelic variants of the rMLST genes of these genomes have been fully catalogued. For species identification purposes, the lowest common taxonomic node (LCTN) of each rMLST allele is calculated based upon the species annotations of the genomes that have that allele. For example, an allele observed in multiple *Streptococcus* species is assigned an LCTN of *Streptococcus* (a genus node), whereas an allele only observed in pneumococcal genomes is assigned an LCTN of *S pneumoniae*.

The rMLST species identification process includes three stages. Firstly, the query genome is scanned against the rMLST allele library using BLASTN (v. 2.12.0) and exact allelic matches are recorded [45]. Secondly, the LCTNs of the matched alleles are mapped onto the nodes of the bacterial taxonomic tree and the lowest observed non-overlapping taxonomic nodes are reported. Finally, the ‘allele support’ of each reported taxonomic node is calculated, which is the number of alleles observed for the reported node divided by the total number of alleles observed across all reported nodes (expressed as a percentage). A reported species node with an allele support above 90% indicates a high degree of confidence in that result.

### Genome quality control

The quality of each genome sequence was assessed based upon several criteria. Firstly, rMLST species identification was applied to each genome and interpreted as pass (only *S pneumoniae* detected, support ≥90%), warning (only *S pneumoniae* detected, support <90%), or fail (*S pneumoniae* not detected or *S pneumoniae* plus other organisms detected). Secondly, genomes were evaluated for evidence of mixed MLST and rMLST alleles: the presence of ≥1 allele at any MLST or rMLST locus (excluding putatively paralogous genes BACT000014 and BACT000062) flagged genomes that were potentially contaminated with non-pneumococcal DNA or consisted of multiple pneumococcal strains.

Thirdly, the interquartile deviation method was used to develop data-derived thresholds for genome size, GC content, number of contigs, N_50_, number of Ns, and number of gaps. Minimum and maximum thresholds (T) were set using the following equations, using *c* = 1.5 for ‘soft’ thresholds and *c* = 2.2 for ‘hard’ thresholds:

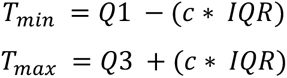

For each metric, genomes were categorised as ‘pass’ (between the soft thresholds), ‘warning’ (between the hard and soft thresholds), or ‘fail’ (outside the hard thresholds). The results of the rMLST species identification, MLST/rMLST allele curation, and genome quality thresholds were implemented in the *S pneumoniae* PubMLST database to enable users to filter PGL data based on genome quality. Finally, genome completeness was assessed using BUSCO (Benchmarking Universal Single-Copy Orthologs) (v.5.4.3) and the lactobacillales_odb10 lineage dataset [46].

### Definition of a cgMLST genotyping scheme

Seventy-one complete pneumococcal genomes were available for download from NCBI in March 2019, of which 29 were excluded from analyses (non-RefSeq genomes, n = 8; genomes with gaps, n = 5; and genomes of STs represented more than once, n = 16, i.e. only one representative of each ST was selected). The chewBBACA software suite (v2.0.16) was run on the remaining 42 genomes using default parameters to create and validate a cgMLST scheme [47]. CreateSchema identified 3,139 complete, non-redundant coding sequences in each genome and alleles were defined for each of these genes using AlleleCall. Putatively paralogous genes (n = 42) were removed using RemoveGenes and annotations for each gene were retrieved using UniprotFinder. Among the remaining 3,095 genes, 1,385 (44.8%) were detected in >95% of reference genomes, excluding genes with alleles that were 20% larger or smaller than the modal length of the distribution of matched genes.

The cgMLST scheme was assessed using 8,263 genomes from an early subset of the PGL, of which 86 were excluded from further analyses: likely contaminated based on presence of multiple alleles at rMLST loci (*n* = 74); or overall size +/− 2 standard deviations from the mean pneumococcal genome size (*n* = 11); or not a pneumococcus, based on rMLST species identification; or seven novel MLST alleles (*n* = 1). The chewBBACA AlleleCall command was re-run on the 8,177 genomes using the 1,385 gene scheme. A further 629 genomes were removed due to high numbers of missing genes (>33) and or warnings (>26). One duplicated genome was removed. The remaining 7,547 genomes comprised the cgMLST scheme validation dataset. AlleleCall flagged 242 genes for manual review and 25 of these were removed from the scheme for these reasons: frequent absence (*n* = 15); alleles 20% smaller than the length mode of the distribution (*n* = 6); transposases (*n* = 2); paralogous hits (*n* = 1); or frequently at the end of assembly contigs (*n* = 1). Finally, a provisional 1,360 gene cgMLST scheme was implemented in PubMLST.

Initial seed alleles were identified using the validation set of 8,177 genomes plus an additional 796 recently assembled genomes. chewBBACA AlleleCall was used to identify alleles and only those at the mode length for each gene were retained. Automated curation within PubMLST (using the BIGSdb scannew.pl script, and thresholds of 98% sequence identity and 98% alignment length to existing alleles) was performed on the 7,547 genomes in the validation dataset. For genomes where alleles/genes were not identified, manual BLASTn searches were performed using lower thresholds of >=70% sequence identity and at least 50% gene length. Truncated alleles, and gene sequences containing insertion sequences, mobile genetic elements or that were otherwise interrupted, were not assigned allele numbers. There were 138 core genes that contained a high proportion of disrupted or missing alleles and were thus excluded since they would be poorly reproducible in a typing scheme. Ultimately, 1,222 core genes were included in the final cgMLST scheme.

To identify the core gene alleles within each genome, three rounds of autocuration of the 1,222 genes were performed within PubMLST on the entire PGL, using the BIGSdb autotag.pl script, which used a subset of alleles as exemplars (defined using find_exemplars.pl for each gene such that all known alleles were within 3% sequence identity to an exemplar allele of the same length). Initially, only hits with >=99% sequence identity and 100% sequence alignment to an exemplar allele were allowed. The autocuration was then repeated, but the sequence identity threshold was first lowered to 97% and then to 96%; the results of these autocuration scans led to a final threshold of 97% sequence identity to exemplar alleles to define new alleles for any new genomes added to the PGL. Manual inspection and curation of 216 genes was also performed as described above to define any alleles not assigned automatically, using BLASTn with a threshold of 95% sequence identity to any known allele. EggNOG-mapper (v5.0.0) was used for functional annotation of cgMLST genes using one randomly selected representative of each coding sequence [48].

### Core genome alignment and phylogenetic analyses

A subset of the PGL was selected for phylogenetic analyses and comprised 1,870 pneumococcal genomes chosen at random, plus nine genomes of ST344 and 21 genomes of ST448. Fifty *S pseudopneumoniae* genomes were included as an outgroup, chosen at random from a curated set of 77 *S pseudopneumoniae* genomes from the rMLST database. The core gene sequences were retrieved by allele number from the sequence definitions database in PubMLST for each of the 1,222 genes, aligned using MAFFT (v7.508; missing alleles were treated as gaps), and concatenated (total length 1.16 Mb) using a custom script [49–50]. A phylogenetic tree was created with FastTree (v2.1.11), using the generalised time-reversible (GTR) model of nucleotide evolution and a single rate for each site (the ‘CAT’ approximation) [51]. This tree was reconstructed to account for recombination using ClonalFrameML [52]. This process was repeated for all lineage-specific phylogenetic analyses. The resulting phylogenetic trees were visualised using ggtree (v3.4.2) and cophenetic distance was calculated using the cophenetic function in the R stats package (v4.2.2) [53].

### Identification of population-wide variation in allelic mismatches

Pairwise allelic mismatch dissimilarities and the Silhouette index (S_t_) were assessed using MSTclust (v0.21b), using a subset of 5000 PGL genomes (which provided detailed analyses in a reasonable processing time) and default parameters [27]. The Silhouette index (S_t_), a measure of cluster cohesiveness, was calculated for the full range of pairwise allelic mismatches among the core genes. The adjusted Rand Index *R_t_* was calculated for each classification level and each pre-existing metric using the adjustedRandIndex function implemented in the mclust R package (v6.0.0) [54].

### Comparison of clustering methods

Clonal complexes based on the seven locus MLST scheme were assigned for all PGL isolates using Phyloviz and named after the predicted founder sequence type(s) [55]. Global Pneumococcal Sequence Clusters (GPSCs) were assigned using PopPunk (v2.6.0) and the GPS reference database (v6) [11, 56]. Mandrake was used to cluster a concatenated alignment of the 1,222 cgMLST genes from a randomly selected subset of 5000 PGL genomes, using default parameters [12].

## Results

### A diverse global dataset of 30,976 published pneumococcal genomes

A total of 211 articles published in 78 journals between 2000-2020 that contained pneumococcal whole genome sequence data were identified (Figure 1a; Supplementary Figure 1). Just over half (n=115, 54.5%) of the publications provided access to all analysed genome data, including reference genomes and contextual isolates. The remaining articles either did not provide the entire genome dataset reported in the publication or there were data integrity issues. This included six publications that contained suppressed genomes, with no published corrections or clarifications regarding the reason for suppression or potential impact on published analyses. Some data issues were resolved by further investigation and/or contacting the corresponding authors for more information (Supplementary Data 1).

**Figure 1:**
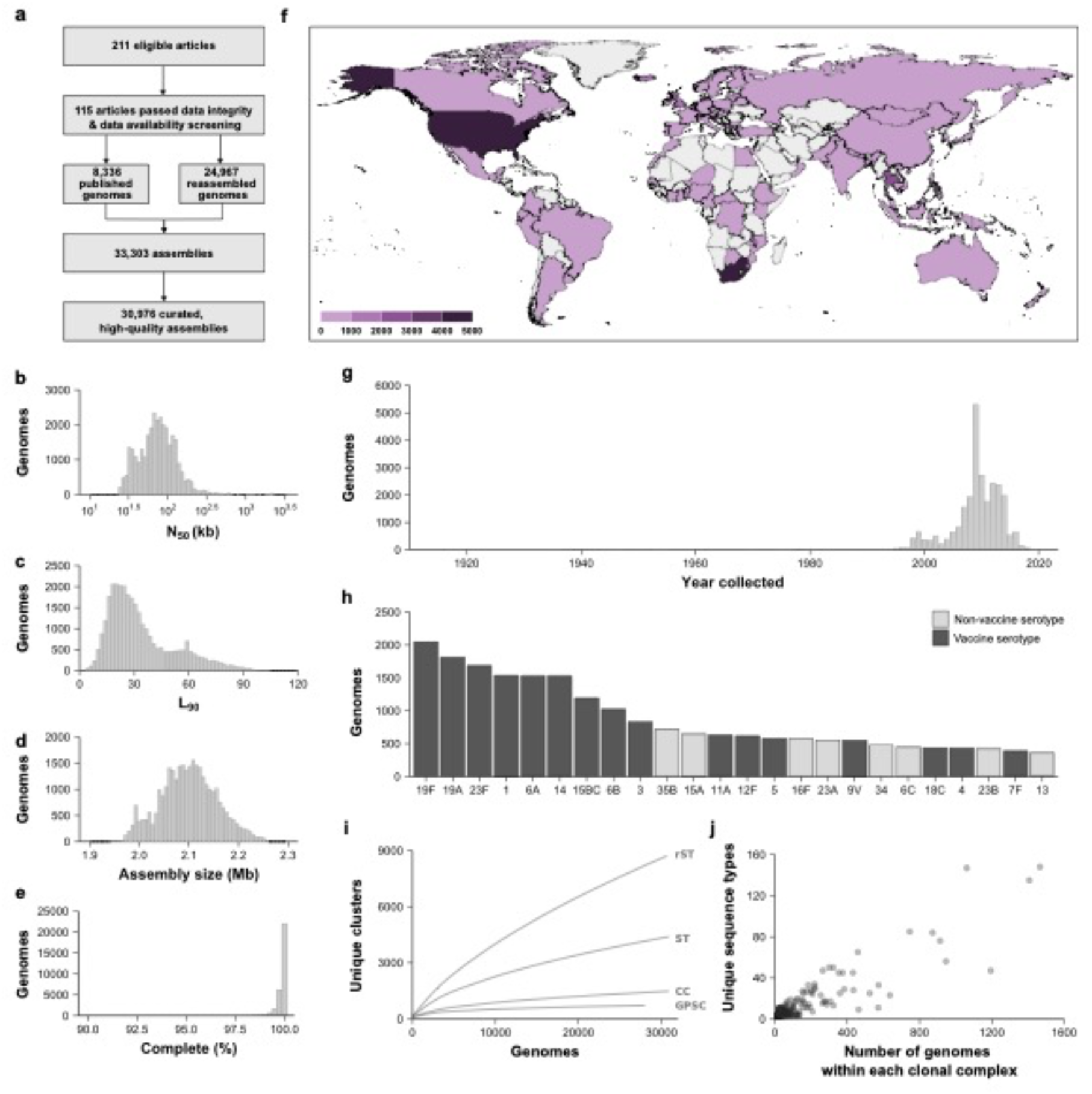
Summary of the creation and contents of the Pneumococcal Genome Library (PGL). a) Schematic overview of the creation of the PGL, containing 30,976 genome assemblies. b-d) Assembly statistics for all genomes. e) Number of complete and single-copy Benchmarking Universal Single-Copy Orthologs for all genomes. f) Worldwide sampling density of the PGL, coloured by the number of genomes from each country. g) Years of collection of the PGL pneumococci. h) The 24 most prevalent serotypes in the PGL, marked to indicate serotypes included in any licensed pneumococcal vaccine. i) Rarefaction curves depicting the number of unique clusters observed for each of the four different taxonomic groups, plotted from random subsets of each sample size in triplicate. Ribosomal sequence type (rST), sequence type (ST; 7-locus multilocus sequence type), clonal complex (CC), and Global Pneumococcal Sequence Cluster (GPSC). j) Number of unique sequence types within each clonal complex.

Overall, the PGL was created with 33,303 genomes from 129 publications. Short-read data were downloaded and assembled for 24,967 genomes, and 8,336 assembled genomes were downloaded from public repositories. Overall, 30,976 (93.0%) genomes passed all quality control metrics (Supplementary Data 2). The genomes were generally highly contiguous (median N_50_ = 74.1 Kb; median L_90_ = 28 Kb; Figure 1b,c) and were of the expected size for a pneumococcal genome (median = 2.10 Mb, range = 1.95-2.26 Mb; Figure 1d). Genomes typically resulted in a full representation of the Lactobacillales BUSCOs (median complete and single-copy BUSCOs = 100%; Figure 1e; Supplementary Data 2; [46]). The remaining 2,165 genomes (6.5%) failed one or more quality checks, of which 67 showed evidence of contamination and 10 were not a pneumococcal genome (Supplementary Data 2). They were excluded from further analyses and filtered out of the PGL.

Pneumococci included in the PGL were recovered from 80 countries across six continents, and more than half of the collection was from South Africa (*n* = 4,887), USA (*n* = 4,273), The Netherlands (*n* = 3,511), The Gambia (*n* = 2,859), and Thailand (*n* = 2,305; Figure 1f; Supplementary Data 3). Provenance data such as carriage/disease status and specimen source were available for 89.4% and 86.1% of pneumococci, respectively. Overall, 42.6% of pneumococci were nasopharyngeal carriage samples. Pneumococci were recovered between 1916-2018, and 58.0% were recovered between 2009-2018 (Figure 1g). Ninety-six serotypes were represented in the PGL, including 24 serotypes that are included in any licensed pneumococcal vaccine (Figure 1h).

There were 8,714 ribosomal sequence types (rSTs), and 4,401 sequence types (STs, seven-locus MLST scheme) that clustered into 1,482 CCs (Supplementary Data 4 and 5) represented in the PGL. Variable-length-k-mer clustering identified 717 GPSCs (Supplementary Data 4). Rarefaction analyses showed that the PGL effectively encompassed the known genetic diversity of pneumococcal population clusters as defined by CCs and GPSCs (Figure 1i) but had a more limited representation of the entire known genotyping diversity as defined by pneumococcal STs and rSTs (>18,000 and >16,000, respectively; Supplementary Data 4). There was a proportional representation of STs across CCs (Figure 1j).

### Defining a pneumococcal core genome multilocus sequence typing scheme

Complete pneumococcal genomes (*n* = 42) were used to define a provisional set of 1,385 non-redundant core genes (coding sequences with start and stop codons) that were present in at least 95% of the 42 complete genomes (Supplementary Data 6). A subset of 7,547 PGL genomes was then used to assess the allele assignments for each of the 1,385 genes. Core genes with high numbers of disrupted or missing alleles were excluded (*n* = 163). This resulted in a final cgMLST v1.0 scheme of 1,222 genes, which was consistent in size with previous estimates of the pneumococcal core genome (range: 912-1,666 genes; [9–10, 59–60]).

The 1,222 core genes were evenly distributed across the pneumococcal genome and had a diverse range of functions (Supplementary Figure 2; Supplementary Data 7). Per-locus analysis using the pairwise homoplasy index found evidence of intragenic recombination in 36.2% of these core genes (*n* = 442*; p* < 4.09 × 10^−5^ after Bonferroni correction; Supplementary Data 8) and greater allelic diversity as compared to non-recombining genes (Supplementary Figure 3).

In total, 634,151 unique core gene alleles (range: 44-2,757 alleles per gene) were assigned across the entire PGL (Supplementary Data 9). 96.3% of the PGL genomes were missing fewer than 25 allele assignments across all 1,222 genes (Supplementary Data 10). The number of core gene alleles assigned per genome did not vary substantially between the majority of CCs, suggesting minimal phylogenetic bias resulting from missing data (Supplementary Figures 4, 5). A core genome sequence type (cgST) was assigned to each pneumococcus that had 25 or fewer missing core gene alleles (*n* = 29,893 genomes), which resulted in 27,531 unique cgSTs (Supplementary Data 4).

For comparison, 50 genomes of *S pseudopneumoniae*, a closely related species, from the rMLST database were screened for the presence of the 1,222 pneumococcal core genes. Between 809 to 923 (median 884.5) of the 1,222 pneumococcal core genes were also identified among the *S pseudopneumoniae* genomes and alleles were assigned to those genes (Supplementary Data 11).

### Defining the structure of pneumococcal populations using the cgMLST scheme

A set of 5000 PGL genomes was chosen at random and used to set population structure boundaries. Pairwise allelic mismatches among the 1,222 core genes formed a discontinuous distribution with three major peaks (Figure 2a; Supplementary Figure 6). The peak centred at 98.8% mismatches exclusively represented pairwise relationships between pneumococci and *S pseudopneumoniae*. The peak centred at 93.0% mismatches predominantly represented comparisons between nontypable pneumococci of either CC344 or CC448 and other PGL pneumococci. CC344 and CC448 were previously implicated in conjunctivitis outbreaks and demonstrated a phylogenetic cluster distinct from other pneumococci [61–62]. The peak centred at 87.9% mismatches represented core gene differences between pneumococci of different CCs, sampled from different countries, in different sampling years. Therefore, a high-level species classification boundary of 96.6% allelic mismatches among 1,180 core genes was used to separate pneumococci from *S pseudopneumoniae*.

**Figure 2:**
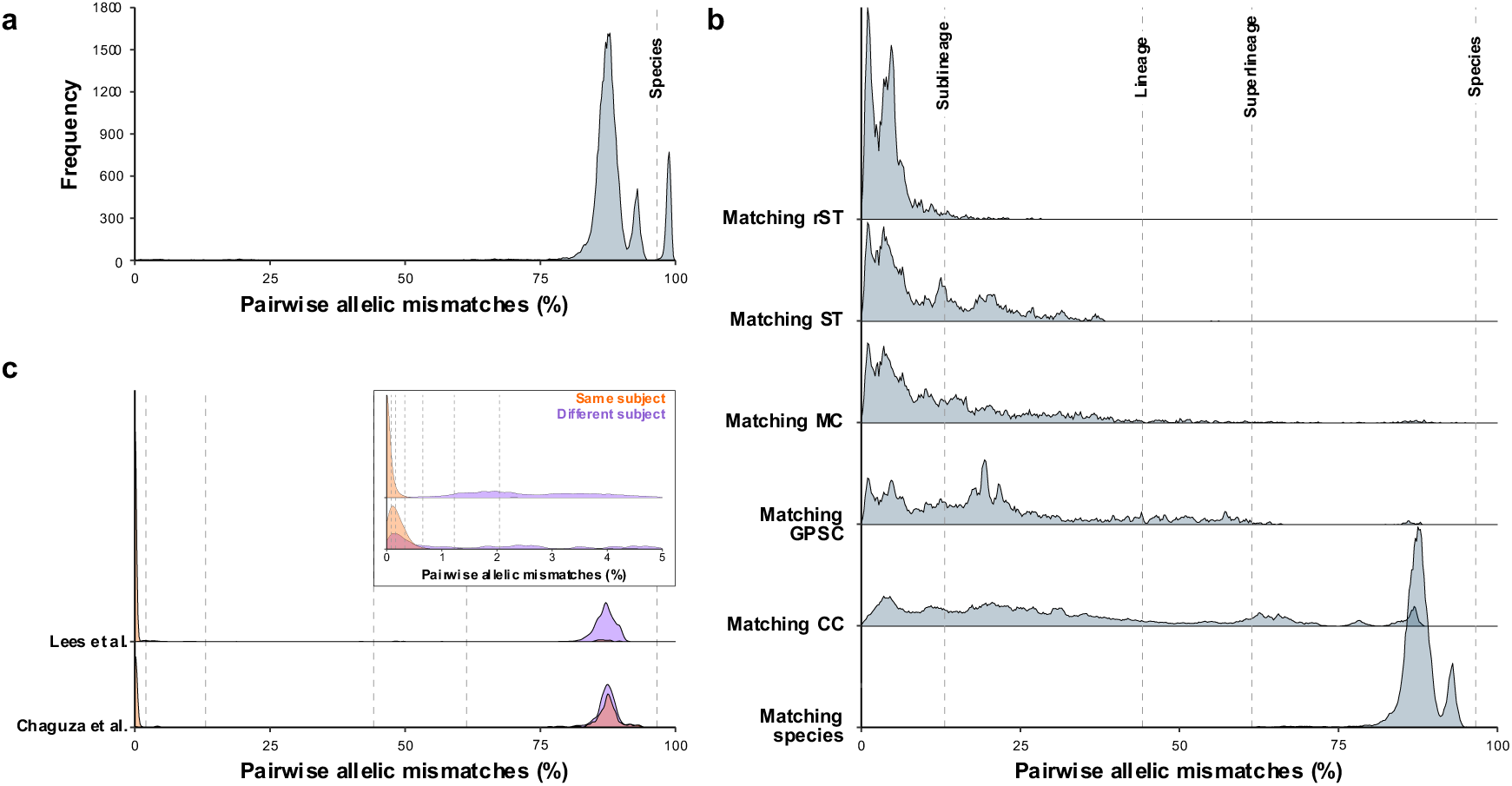
Distribution of pairwise core genome MLST (cgMLST) allelic differences across globally dispersed pneumococcal populations. a) Pairwise cgMLST allelic differences between 5000 randomly selected PGL genomes. X-axis values are plotted as a percentage of all 1,222 cgMLST loci, excluding pairwise relationships where one or both loci had unassigned alleles. The species boundary separates pneumococcal and *S pseudopneumoniae* genomes. b) Distribution of pairwise cgMLST allelic differences between genomes belonging to the same taxonomic group: ribosomal sequence type (rST), sequence type (ST; 7-locus multilocus sequence type), Mandrake cluster (MC), Global Pneumococcal Sequence Cluster (GPSC), clonal complex (CC), and bacterial species. c). Distribution of pairwise cgMLST allelic differences between pneumococci isolated from the same individual either concurrently (blood and cerebrospinal fluid samples) or longitudinally (1-52 weeks post-birth), as published by Lees et al and Chaguza et al, respectively [64–65].

There was no clear peak structure among pneumococci with fewer than 80% pairwise mismatches (Figure 2a; Supplementary Figure 6). To define lineage boundaries within the data, the distributions of allelic mismatches within pneumococcal groups that were defined by ribosomal MLST, seven-locus MLST, Mandrake clusters, GPSCs, and CCs, were compared (Figure 2b). In all cases the distributions were positively skewed and the majority of pairwise relationships were found below 50% mismatches.

Just over 98% of pneumococci within the same GPSC or Mandrake cluster, and 85% of pneumococci within the same CC, had 61.4% (n=750 genes) or fewer core gene allelic mismatches, thus a ‘superlineage’ boundary was defined at 61.4% mismatches. Core gene allelic mismatches among pneumococci with the same ST ranged from 0.0-38.2% but were predominantly low (median = 6.6% core gene allelic mismatches); therefore, a ‘lineage’ boundary (44.2% allelic mismatches or 540 genes) was defined to encompass all matching STs, which coincided with the highest S_t_ value.

Ribosomal STs (rSTs) catalogue sequence variation at ribosomal protein genes, which are conserved, robust to recombination effects, and typically used to differentiate bacterial species [18, 63]. The PGL data demonstrated that pneumococci with matching rSTs also shared the majority of core gene alleles (97.9% of pneumococci with matching rSTs had fewer than 13.1% (n=160 genes) core gene allelic mismatches). A similar observation was made among STs (69.3% of pneumococci of the same ST had fewer than 13.1% core gene allelic mismatches); therefore, we defined a ‘sublineage’ boundary at 13.1% core gene mismatches.

Finally, to differentiate very closely-related pneumococci more precisely, additional boundaries at 2.1%, 1.2%, 0.7%, 0.3%, 0.16% and 0.1% (corresponding to 25, 15, 8, 4, 2, and 1 gene(s), respectively) were used. The 2.1% boundary defined ‘clonal group’ and the remaining boundaries defined ‘clonal subgroups’. Finally, a zero-mismatch boundary grouped cgMLST profiles that differed only by missing data.

To validate these boundaries, cgMLST allelic variation between pneumococci recovered from the same individuals either sampled concurrently (blood and cerebrospinal fluid; [64]), or longitudinally (1-52 weeks from birth; [65]), were compared (Figure 2c). Pneumococci recovered from the same study subjects formed clear peaks at <1% core gene allelic mismatches (Figure 2c).

### Multilevel clustering of PGL genomes

There were 27,531 unique core genome sequence types (cgSTs) in the entire PGL. Multilevel single-linkage clustering was performed using one representative of each unique cgST, which iteratively clustered cgMLST profiles based on pairwise allelic mismatches, and cgST profiles below each mismatch threshold were assigned to the same cgST cluster. Clustering was performed sequentially, with the input order determined by the tip order of cgSTs in a minimum spanning tree that was constructed based on allelic profile similarity.

During cgST clustering, a cgMLST-based life identification number (cgLIN) code was also assigned to each cgST. This multi-positional barcode has 11 numbers that reflect the cgST cluster assignment at its respective threshold at the boundaries defined above, ie species, superlineage, lineage, sublineage, clonal group, clonal subgroups, and the zero threshold. When any two barcodes were compared, the leftmost point of numerical dissimilarity reflected the threshold at which genomes no longer clustered together (Figure 3a). Therefore, cgLIN barcodes reflected an approximation of the phylogenetic relationships between genomes, for example: superlineage 0_10 was composed of four lineages; lineage 0_10_0 was divided into five sublineages; and sublineage 0_10_0_24 subdivided into seven distinct clonal groups (Figure 3b).

**Figure 3:**
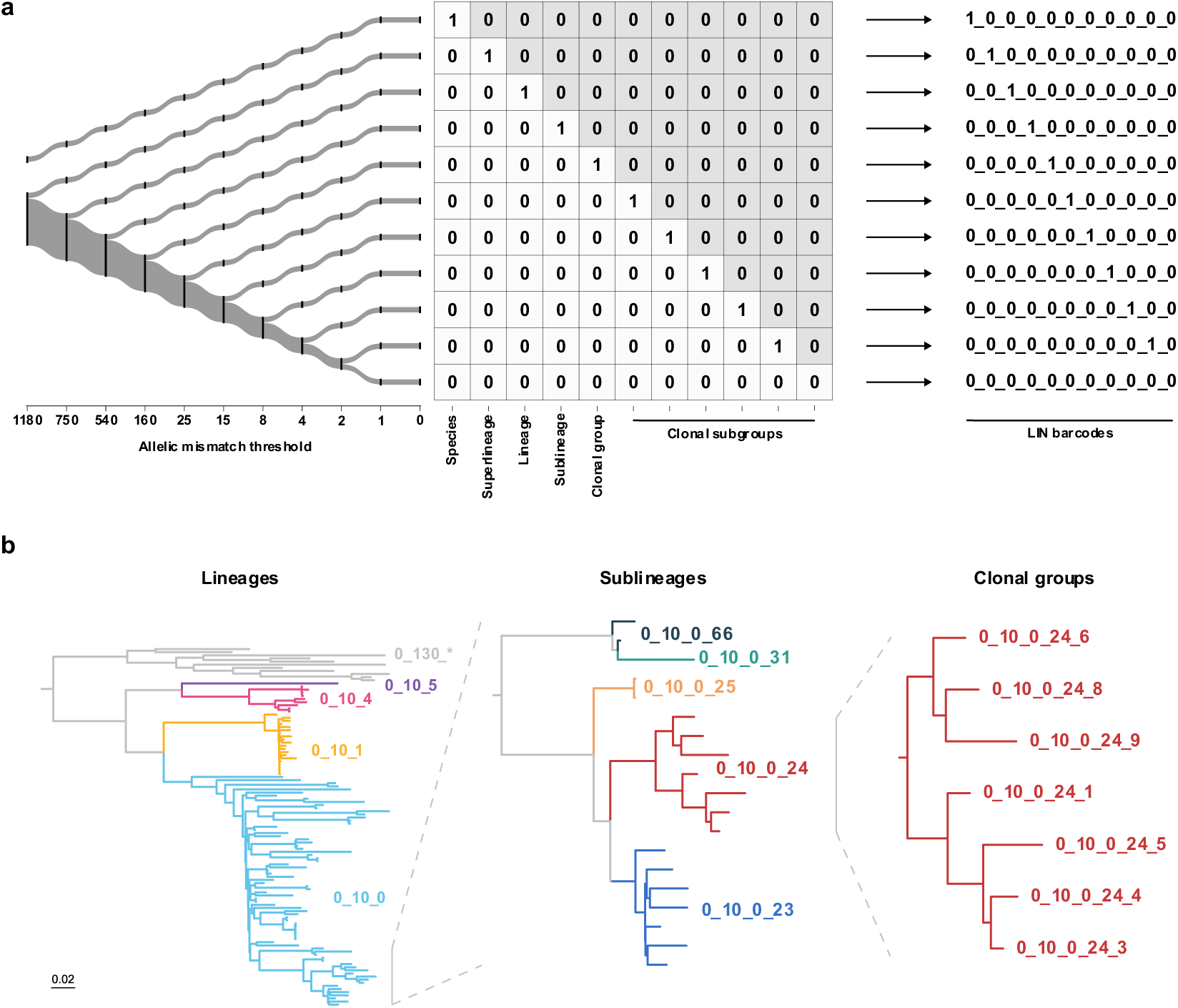
Schematic representation of the cgMLST life identification number (LIN) barcoding approach. a) Overview of LIN barcode construction for each taxonomic level. A simplified phylogeny (left) depicts the allelic mismatch values used to define each single-linkage clustering threshold. The leftmost point of numerical dissimilarity in the barcode indicates the threshold at which genomes no longer cluster together. b) A demonstration of phylogenetic relationships using superlineage 0_10 as an example and superlineage 0_130 as an outgroup. The relationships among lineages (left), sublineages (middle) and clonal groups (right) and the corresponding LIN barcodes are shown.

In total, 27,524 unique cgLIN codes were defined for 29,895 PGL genomes (96.5% of all PGL genomes; Supplementary Data 4). cgLIN codes defined 407 superlineages, 726 lineages, 2,782 sublineages and 12,246 clonal groups within the PGL, and 71.8% of the lineages and 50.5% of the sublineages were represented by >1 genome (median: 2 genomes; range: 1-1,298; Supplementary Data 12). Over half of the PGL was represented by 15 superlineages and 24 lineages.

A subset of 1,800 pneumococcal and 50 *S pseudopneumoniae* genomes was chosen at random from the PGL and rMLST databases, respectively, to determine the phylogenetic congruence of the assigned lineages. A maximum-likelihood phylogeny was reconstructed using nucleotide sequence alignments of all 1,222 cgMLST genes (Figure 4a). The phylogeny identified 221 lineages, and the 20 most prevalent lineages (47.4% of PGL genomes) were predominantly monophyletic and formed discrete phylogroups (Supplementary Figure 7). Comparison of allelic dissimilarity and core genome divergence (represented by cophenetic distance of the phylogeny) demonstrated only limited overlap of classification levels (Figure 4b).

**Figure 4:**
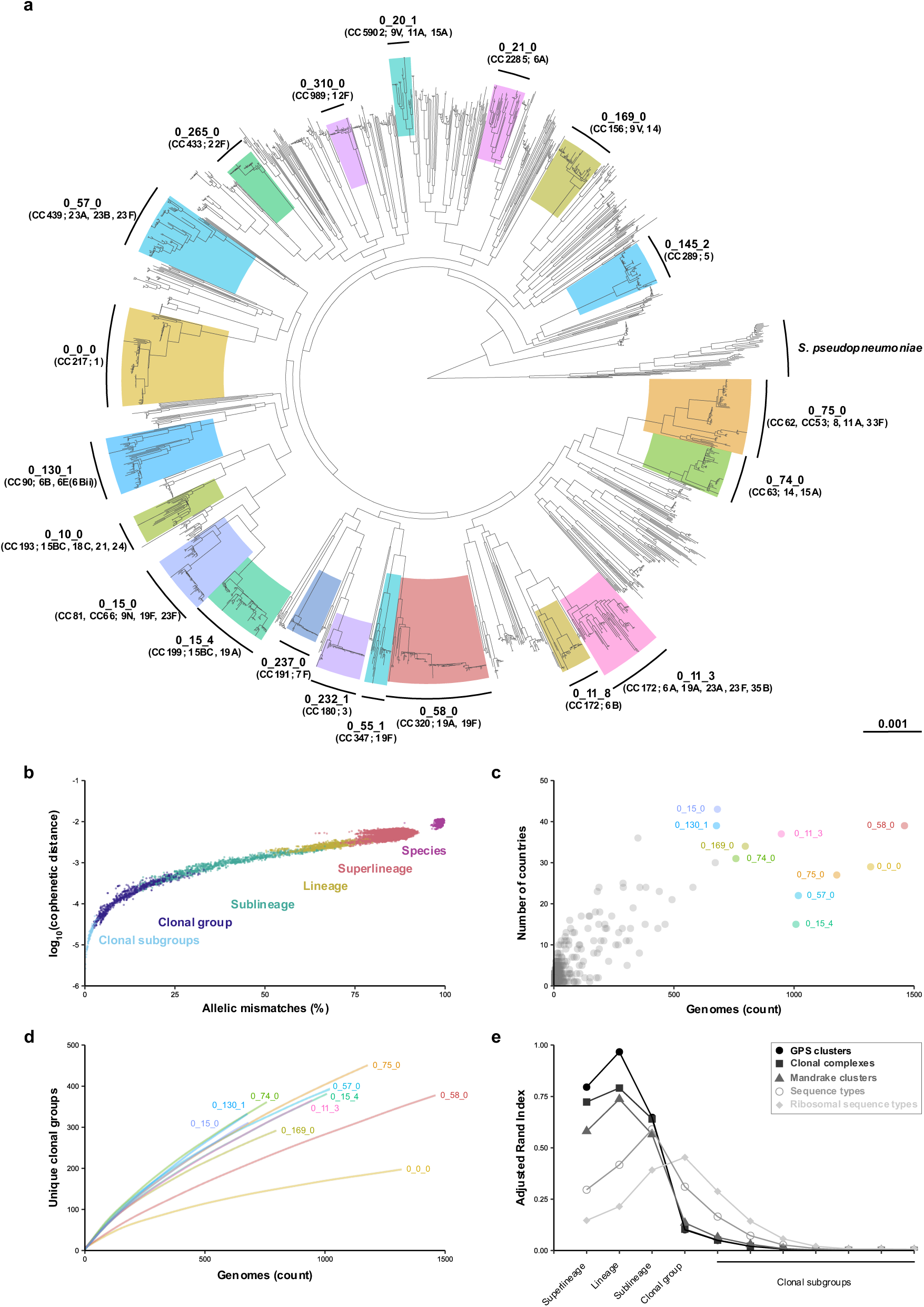
Global pneumococcal population structure and taxonomic classification. a) Maximum-likelihood phylogeny of 1,800 pneumococcal and 50 *S pseudopneumoniae* genomes, based on a nucleotide alignment of 1,222 cgMLST loci and rooted with *S pseudopneumoniae*. The 20 most prevalent pneumococcal lineages are highlighted and annotated with CC, last three differentiating digits of the LIN barcode, and predominant serotype(s). b) Comparison of similarity among genomes represented by the phylogeny in panel a. Pairwise relationships were calculated for all 1,850 genomes and randomly subset down to 50,000 pairs. Points are coloured by the closest inferred taxonomic relationship. c) Geographical diversity of pneumococcal lineages. The ten most prevalent lineages are highlighted. d) Rarefaction curves depicting the number of unique clonal groups observed for each of the ten most prevalent lineages, plotted from random subsets of each sample size in triplicate. e) Similarity between cgMLST cluster classification levels defined in this publication and other measures of population structure. Concordance between clustering metrics was measured using the adjusted Rand Index.

Among lineages, 45.5% were comprised of pneumococci from more than one country, and 30.0% were from more than one continent (Supplementary Data 12). Consistent with previous studies, individual lineages were normally associated with one or a small number of serotypes (mode number of serotypes per lineage, 1; range, 1-21 serotypes; Supplementary Figure 8, Supplementary Data 12,13). Rarefaction analysis suggested wide variability in substructure diversity within each lineage (Figure 4d). For example, lineage 0_0_0 (serotype 1, CC217) demonstrated low clonal subgroup diversity despite extensive sampling (*n* = 1,319 genomes), as compared to other lineages such as 0_15_0 (serotype 23F, CC81 and CC88; *n =* 681) and 0_75_0 (serotype 11A, CC53 and CC62; *n* = 1178). The concordance between cgLIN clustering and other approaches was calculated using the adjusted Rand index (ARI), a measure of similarity between clustering approaches. At the lineage level, cgLIN clustering was nearly identical to GPSCs (ARI = 0.97), and highly concordant with clonal complexes (ARI = 0.79) and Mandrake clusters (ARI = 0.74) (Figure 4e).

## Discussion

Data sharing is essential for reproducibility and transparency in science. The open data movement has gained momentum, and many publishers, funders, and organisations encourage or require authors to share data [66]. There are strong arguments for data sharing in the field of pathogen genomics, particularly in a public health context such as when managing disease outbreaks [3, 67–68]. The COVID pandemic highlighted the benefits of rapid data sharing since publicly available SARS-CoV-2 genome data informed the development of vaccines, infection control strategies, and diagnostic assays [69]. An open data culture allows datasets to be repurposed to advance our understanding of the epidemiology, evolution, and biology of important human pathogens.

The PGL is a comprehensive, open-access database of nearly 31,000 curated pneumococcal genomes from a broad range of countries, serotypes, genotypes, clinical manifestations, and sampling years. The genomes and contextual data (provenance and phenotype) are easily accessible through the web-based PubMLST platform, which provides an extensive suite of third-party software to facilitate downstream analyses. The PGL aggregates existing genomes and helps to highlight underrepresented geographical regions that should be the focus of future genomic surveillance efforts.

Although the PGL contained genome data from 129 publications, many publications had data integrity issues that could not be resolved by the time of publication. This is contrary to open data principles and policies and is a barrier to the reproduction of published analyses to assess the validity of research findings [70–74]. Importantly, it also precludes integration of these datasets into surveillance efforts, potentially leading to unnecessary duplication of efforts or distorting estimates of global coverage of pneumococcal genomic surveillance. More stringent checks of adherence to open data standards during the publication process are necessary and could be made easier by the standardisation of data input formats. It also seems unlikely that these issues are unique to pneumococcal genomes and an assessment of published genome datasets for other microbial species is warranted.

Compared to the original seven-gene MLST, cgMLST provides much higher resolution of pneumococcal population structure, comparable to that of single nucleotide variants or variable length k-mer comparisons. Moreover, by retaining the methodological approach and increasing the number of genes, cgMLST retains the advantages of MLST, namely: consistency; standardised nomenclature; rapid allele assignment; and the representation of alleles as numerical indices [16, 25, 75]. Furthermore, this cgMLST scheme is differentiated from other typing schemes by extensive manual curation of genes that were identified in a large validation set of genomes, which led to a large set of stable, reliably sequenced core genes and a robust genotyping scheme. Additionally, by applying this cgMLST scheme to the large PGL dataset we have already created an extensive database of pneumococcal allelic variation, and this reduces the amount of curation required going forward as new genomes are added to the PGL.

Using a wide range of fixed clustering thresholds, the pneumococcal population was differentiated at varying phylogenetic and epidemiologically relevant scales, which provided greater resolution than typing schemes with single clustering levels. The added complexity of multilevel clustering was counterbalanced with an intuitive barcoding system and classification level, to provide a consistent description of each clustering level. These analyses revealed many pneumococcal lineages, but since there were few obvious discontinuities in the percentage of pairwise allelic mismatches across the PGL, existing clustering metrics were used to guide the definition of LIN boundaries. The observed flat structure in pairwise allelic mismatches might be explained by geographical, serological or genotype-specific population substructures that are obscured by the large size of the PGL, or might be due to the naturally high recombination rates of pneumococci that may have created a natural gradient of allelic similarity. The analyses of clonal groups and clonal subgroups aimed to differentiate very closely related pneumococci and those boundaries were validated with studies of multiple and longitudinal sampling.

In conclusion, we have created a high-quality, open-access genomic resource that is representative of pneumococcal global diversity, and a novel pneumococcal cgMLST scheme and barcoding system to define and evaluate genetic lineages, in a manner that reflects the complex structure of pneumococcal populations.

## Data availability

The PGL, cgMLST scheme, and LIN barcodes are available from PubMLST (https://pubmlst.org/bigsdb?db=pubmlst_spneumoniae_isolates_pgl). Genome accession numbers are available within each isolate record in PubMLST and in Supplementary Data 2.

## Code availability

The code used for data analyses is available at: https://github.com/brueggemann-lab

## Authors and contributors

Conceptualisation: MJJvR, ABB. Literature search and data acquisition: MJJvR, AF. Data curation: MJJvR, DJB, AF, JEB, KAJ, ABB. Genome assembly: JEB. Data analyses: MJJvR, DJB, JEB, KAJ, ABB. PubMLST platform and software development: JEB, KAJ, MCJM. Data visualisation: MJJvR, DJB. PubMLST funding and infrastructure: KAJ, MCJM, ABB. Writing of first draft: MJJvR, DJB, ABB. All authors contributed to the final version of the manuscript.

## Acknowledgements

We gratefully acknowledge all authors who shared their genomic data on publication, thereby making the PGL possible. Mario Ramirez and João Carriço were helpful in the early development of this work, in particular with discussions about chewBBACA, and the provision of SPAdes genome assemblies. Madeleine Butler and Femke Ahlers contributed to data curation in the early development of the cgMLST scheme as part of their bioinformatics training.

