## Supplementary Figures for "Development of the Pneumococcal Genome Library, a core genome multilocus sequence typing scheme, and a taxonomic life identification number barcoding system to investigate and define pneumococcal population structure"

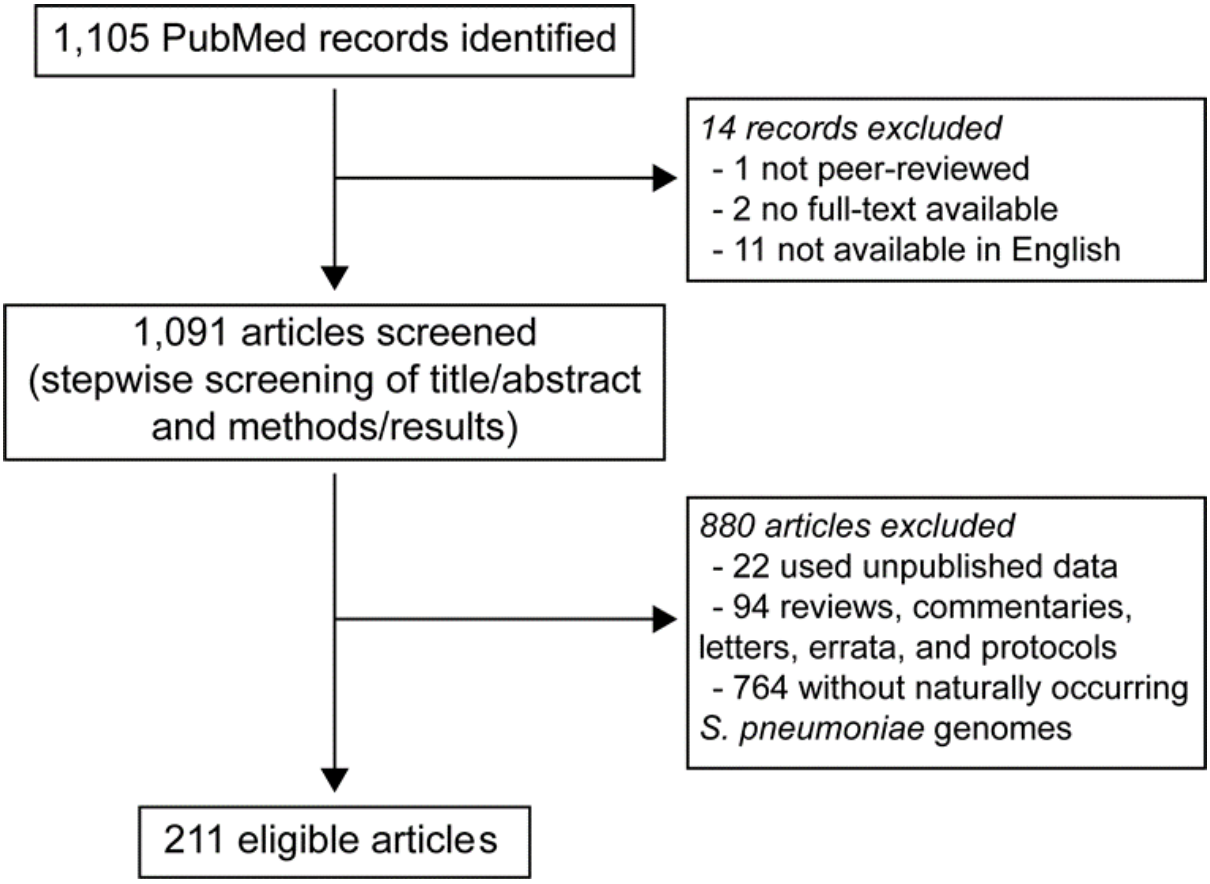


**Supplementary Figure 1. Selection of publications for inclusion in the Pneumococcal Genome Library.**

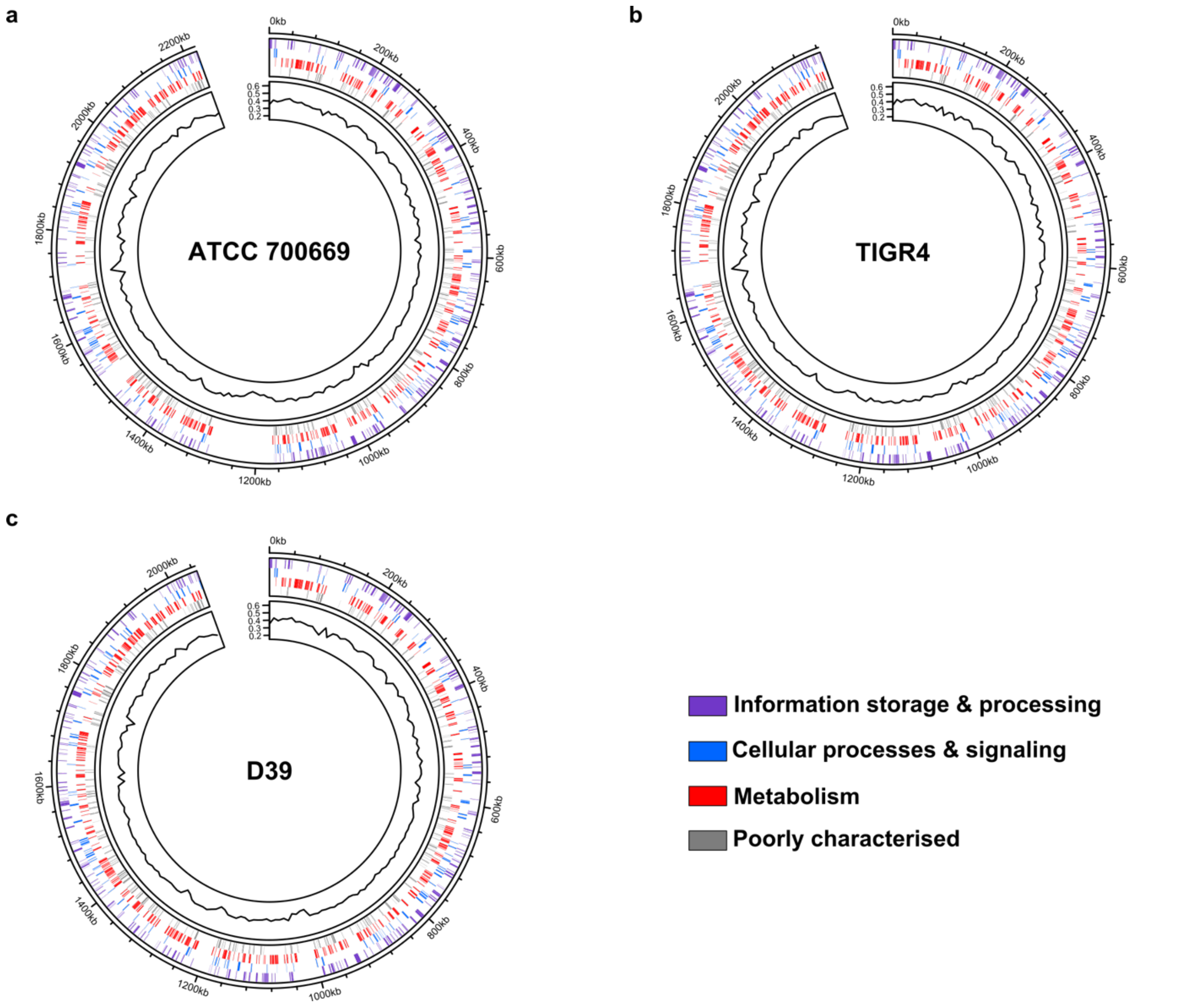


**Supplementary Figure 2: Distribution of cgMLST loci across three closed (complete) pneumococcal reference genomes.** NCBI RefSeq accession numbers for each reference genome are as follows: a) ATCC 700669, GCF_000026665.1; b) TIGR4, GCF_000006885.1; and c) D39, GCF_000014365.2. The outer ring in each plot represents cgMLST loci coloured by predicted gene function and Clusters of Orthologous Groups (COG) category for each gene cluster, as inferred by eggNOG-mapper. Loci without a predicted function were marked as ‘Poorly characterised’. The inner ring depicts the GC content, calculated as the median value of 1 kb windows along each complete genome.

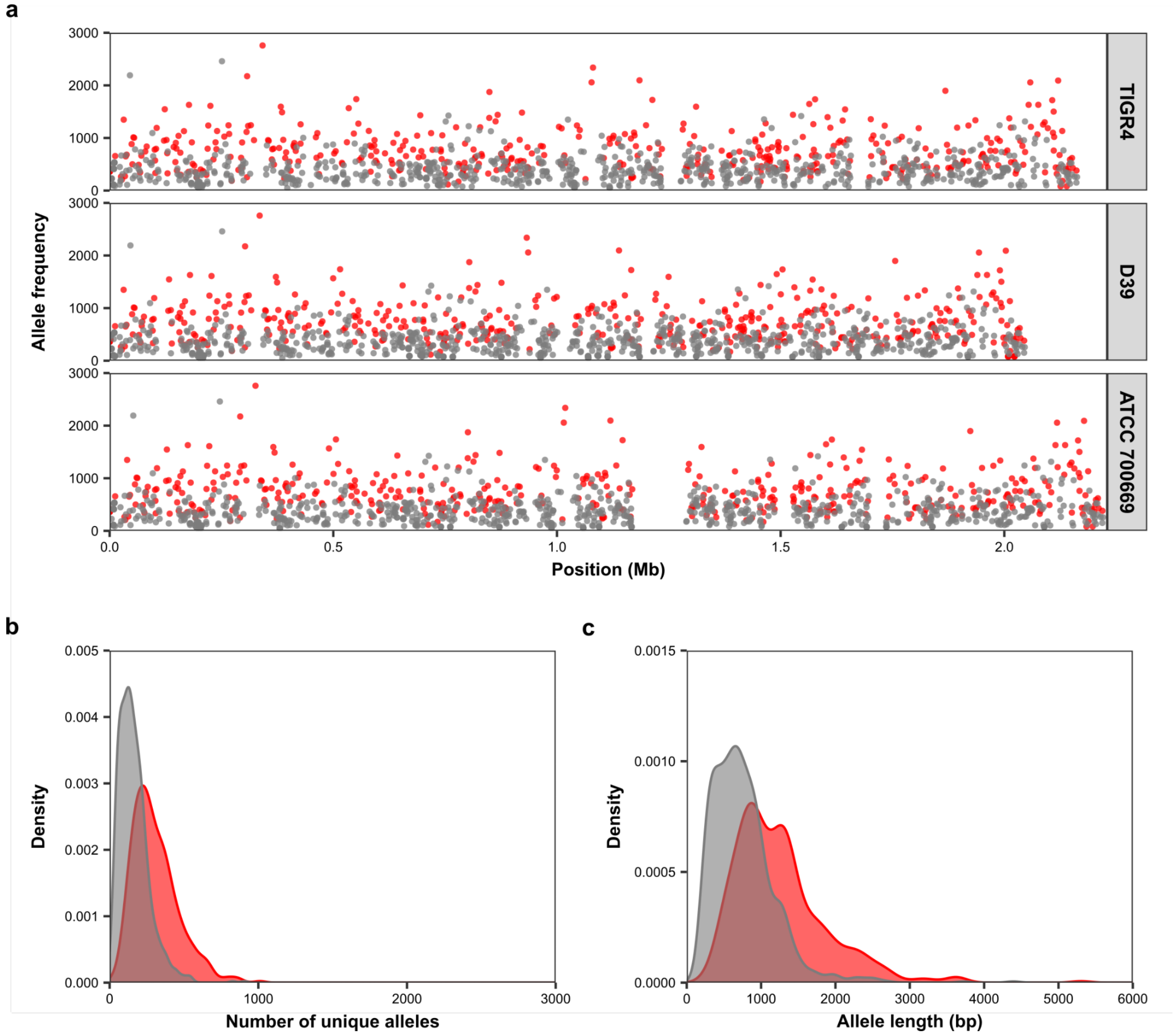


**Supplementary Figure 3: Allele and recombination frequencies of the cgMLST core genes.** Red points and peaks represent loci with evidence of significant intra-gene recombination, as estimated by the pairwise homoplasy index after Bonferroni correction ($p < 4.09 \times 10^{-5}$) compared to those loci without significant recombination (grey). a) Allele frequency at each cgMLST gene across three pneumococcal reference genomes. b) Distribution of total counts of unique alleles for each cgMLST genes ($n$ = 1,222). c) Allele length distribution for each cgMLST gene ($n$ = 1,222).

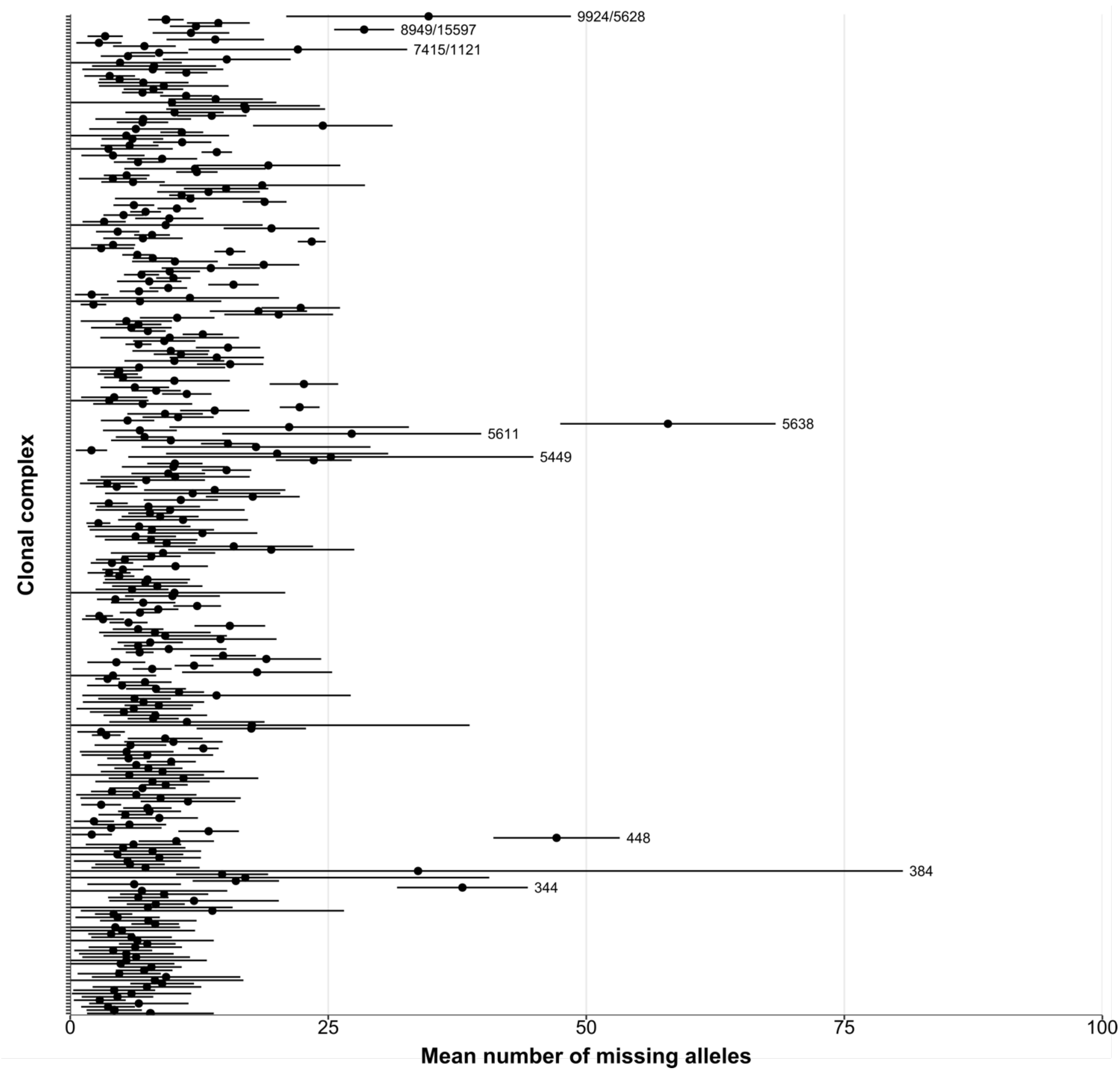


**Supplementary Figure 4: Core gene allele assignments for each clonal complex.** Black points represent the per-genome mean number of cgMLST genes without an allele assignment for each clonal complex, for all clonal complexes with >= 10 genomes. Horizontal lines represent the standard deviation of the mean. Clonal complexes with a mean >25 missing alleles are labelled.

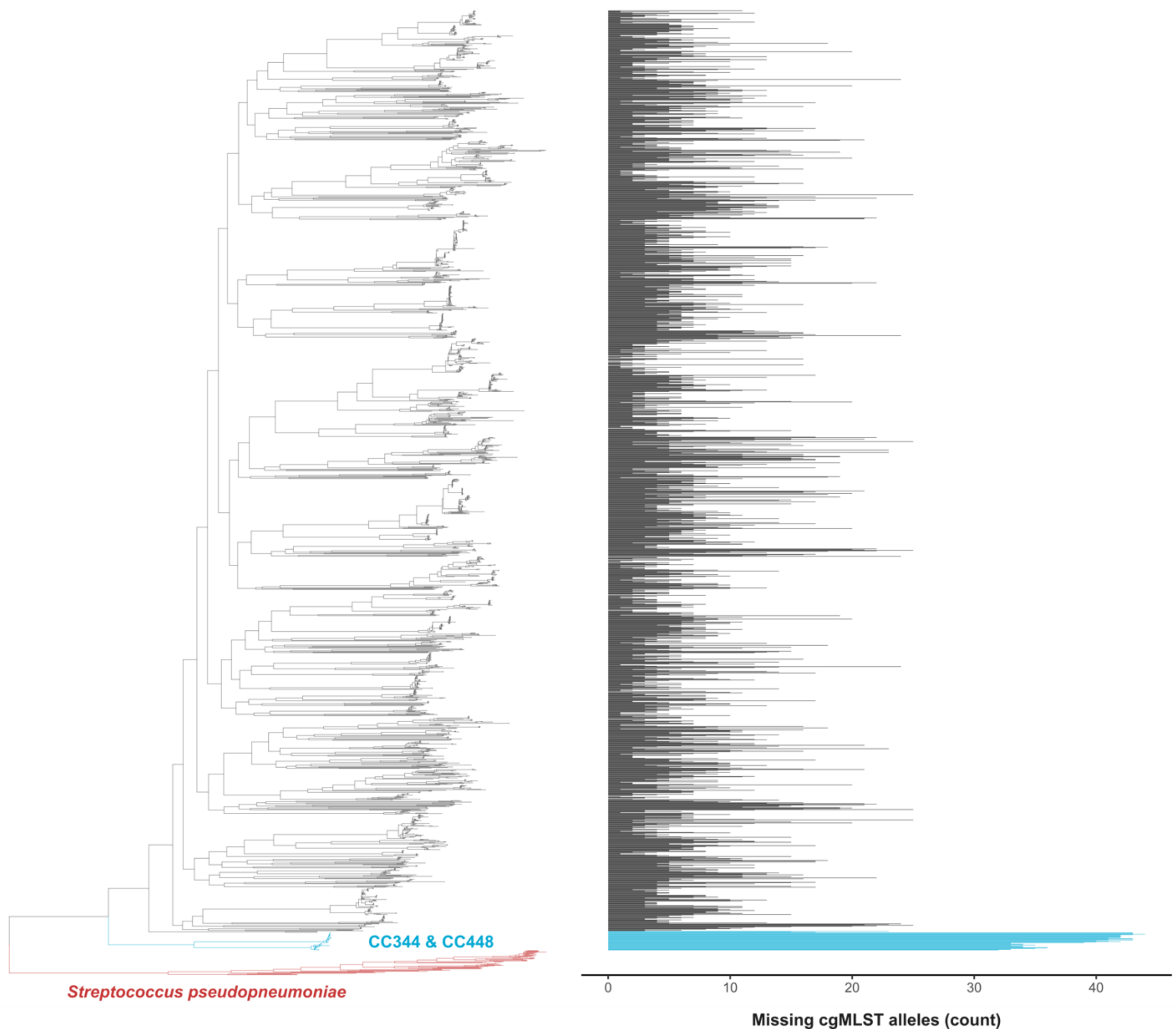


**Supplementary Figure 5:** Left: Maximum-likelihood phylogenetic tree representing data from 1,800 randomly selected PGL genomes plus 50 *S. pseudopneumoniae* genomes (red); representatives of clonal complexes 344 & 448 are highlighted in blue. The phylogenetic tree was constructed using a nucleotide alignment of 1,222 cgMLST loci and rooted with *S pseudopneumoniae*. Right: Per-genome rates of allele missingness (cgMLST genes with unassigned alleles), out of a total of 1,222 genes. Bars representing values of clonal complexes 344 & 448 are highlighted in blue.

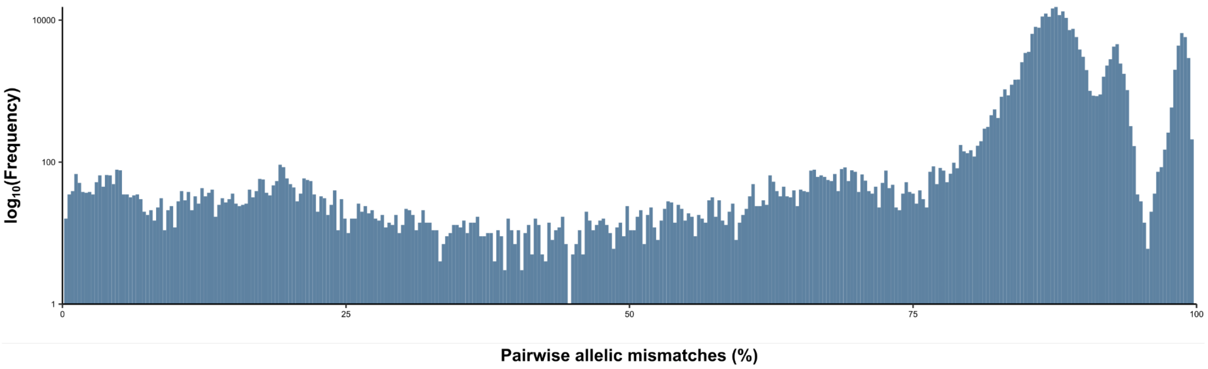


**Supplementary Figure 6:** Distribution of pairwise cgMLST allelic differences between 5,000 randomly selected genomes in the PGL. Allelic mismatches on the x-axis are plotted as a percentage of all 1,222 cgMLST loci, excluding pairwise comparisons where one or both of the pair being compared had an unassigned allele at that locus.

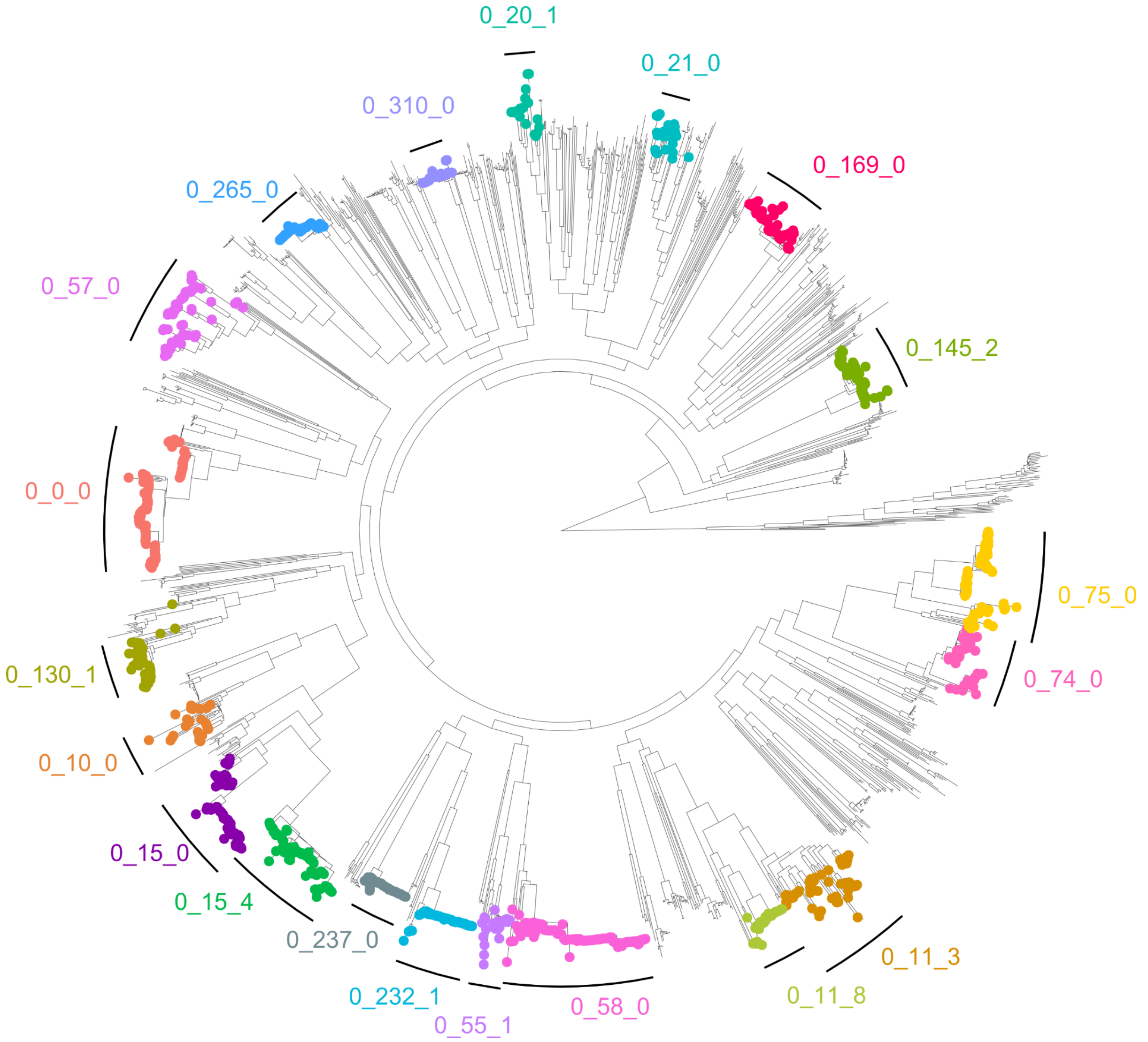


**Supplementary Figure 7: Maximum-likelihood phylogenetic tree representing 1,800 randomly selected PGL genomes plus 50 *S pseudopneumoniae* genomes.** The phylogenetic tree was constructed using a nucleotide alignment of 1,222 cgMLST loci and rooted with *S pseudopneumoniae*. Genomes contained in the 20 most prevalent pneumococcal lineages are shown as coloured tip points, and annotated with the first three LIN barcode identifiers, ie species_superlineage_lineage.

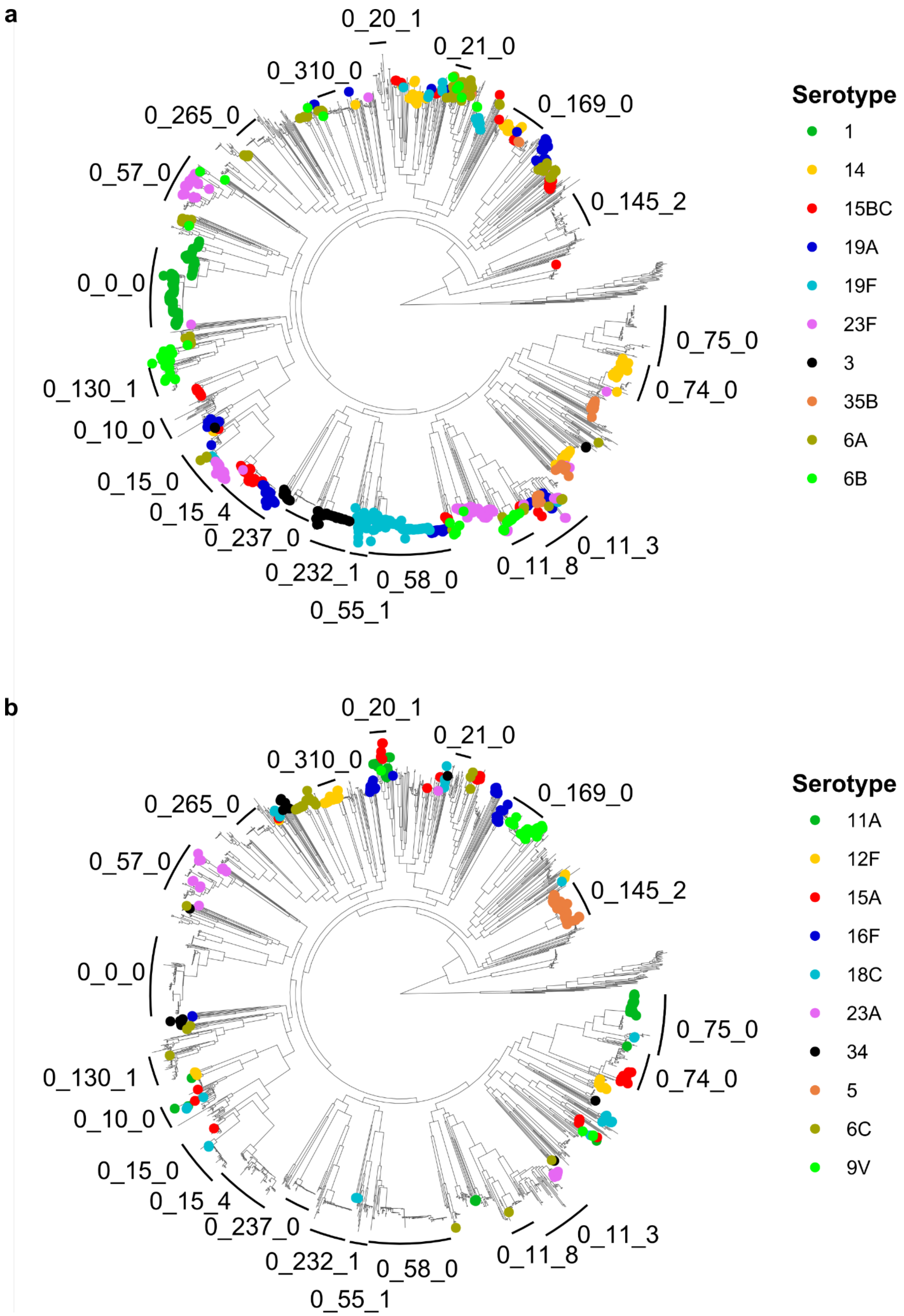


**Supplementary Figure 8: Maximum-likelihood phylogenetic tree representing data from 1,800 randomly selected PGL genomes plus 50 *S pseudopneumoniae* genomes.** The phylogenetic tree was constructed using a nucleotide alignment of 1,222 cgMLST loci and rooted with *S pseudopneumoniae*. Genomes representing the 20 most prevalent serotypes are shown by coloured branch tips.
